# Hyaluronic acid particle hydrogels decrease cerebral atrophy and promote pro-reparative astrocyte/axonal infiltration in the core after ischemic stroke

**DOI:** 10.1101/768291

**Authors:** Elias Sideris, Aaron Yu, Jun Chen, S Thomas Carmichael, Tatiana Segura

**Affiliations:** Department of Chemical and Biomolecular Engineering, University of California Los Angeles, Los Angeles, CA, United States; Departments of Biomedical Engineering, Neurology, and Dermatology, Duke University, Durham, NC, United States; Department of Neurology, David Geffen School of Medicine, University of California Los Angeles, Los Angeles, CA, United States

## Abstract

The death rate due to stroke is decreasing, resulting in more individuals living with stroke related disabilities. Following stroke, dying cells contribute to the large influx of highly reactive astrocytes and pro-inflammatory microglia that release cytokines and lead to a cytotoxic environment that causes further brain damage and prevents endogenous repair. Paradoxically, these same cells also activate pro-repair mechanisms that contribute to endogenous repair and brain plasticity. Here, we show that the direct injection of a hyaluronic acid based microporous annealed particle (HA-MAP) hydrogel into the stroke core reduces the percent of highly reactive astrocytes and increases the percent of alternatively activated microglia in and around the lesion. Further, we show that HA-MAP hydrogel promotes reparative astrocyte infiltration into the lesion, which directly coincides with axonal penetration into the lesion. Additionally, HA-MAP injection decreases cerebral atrophy and preserves nigrostriatal bundles after stroke. This work shows that the injection of a porous scaffold into the stroke core can lead to clinically relevant decrease in cerebral atrophy and modulates the phenotype of astrocytes and microglia towards a pro-repair phenotype.

## Introduction

Stroke is the leading cause of adult long-term disability and affects 795,000 Americans every year ^1^. Strokes occurs in two main forms, hemorrhagic or ischemic. Ischemic strokes are caused by an obstruction within a blood vessel, and account for 87% of all strokes and have a 32% mortality, ^2, 3^. For some stroke survivors (∼90% for ischemic stroke), the injury leaves them with a serious disability. There are currently no FDA approved therapies to treat long-term disability, leaving physical therapy as their only medical treatment.

Immediately following stroke onset, the lack of oxygen and nutrients causes significant cell death, a large influx of microglia/macrophages and the activation of highly reactive astrocytes, which release pro-inflammatory cytokines ^4, 5^, and lead to further neuronal death and a clearance of cellular debris ^6^. Over time, the brain’s defense mechanism is to compartmentalize the injured tissue from the surrounding tissue via an astrocytic and fibrotic scar, preventing repair in the stroke core ^7^. With time the stroke core is devoid of vessels and axons and cerebral atrophy occurs (brain shrinkage). Atrophy in the motor cortex accounts for at least a portion motor deficit in stroke patients^8^ and occurs at a rate of 0.95% of initial volume in the stroke hemisphere ^9^. There are currently no therapies to prevent or treat cerebral atrophy, which is correlated with dementia ^10^, depression ^11^, and reduced motor function ^8^

Astrocytes can both aid and obstruct stroke recovery ^12, 13^, with complete ablation of astrocytes resulting in a worse outcome after stroke^14^. Astrocytes are able to communicate with multiple neurons via secreted and contact-mediated signals, can coordinate the development of synapses ^15-19^ and neural circuits ^20^, yet astrocytes can limit long term repair and regeneration when a pro-inflammatory phenotype is adopted and a scar is formed ^12, 13^. Recently others ^21-23^ and we ^24, 25^ have published a number of publications injecting hydrogels into the stroke cavity. These studies have demonstrated that the stroke core can accept a significant gel injection volume without damage and that these gels can vary widely in composition ranging from hydrogels formed from decellularized native tissue ^26^, to peptide derived hydrogels ^27^, to synthetic hydrogels ^28^. Although biomaterial strategies for brain repair have investigated astrocytes after stroke in the context of the glial scar, no biomaterial has been described that can modulate the astrocyte phenotype from inflammatory to pro-regenerative. Herein, we explore the use of microporous annealed particle (MAP) hydrogels to modulate astrocyte and microglia phenotype towards a pro-repair phenotype and the evaluation of the resulting reparative response.

### Characterization of MAP hydrogel and stroke tissue

Granular hydrogels are materials generated from hydrogel microparticle (HMP) building blocks, generated using a droplet generator flow focusing microfluidic device (**Fig. 1A**). HMPs used in this study are ∼80µm diameter (**Fig. 1B, C**), generated using a hyaluronic acid (HA) backbone, and peptides as crosslinkers (MMP degradable) and ligands for integrin binding (RGD). We crosslink HMPs to each other using the coagulation enzyme factor XIIIa (FXIIIa) to generate a stable scaffold, with a Young’s Modulus of ∼927 Pa (**Fig. 1D**). We termed these linked HMP hydrogels <u>M</u>icroporous <u>A</u>nnealed <u>P</u>article (MAP) scaffolds. MAP scaffolds have micron sized voids (pores) formed in between packed beads (**Fig. 1E**). Others and we have utilized MAP scaffolds for cell culture in vitro ^24, 29-35^ and support tissue ingrowth in vivo^24, 30-32^. HA-MAP scaffolds can be injected into the stroke core at 5-days post wounding without deforming the recipient hemisphere (**Fig. 1F, G**). HA-MAP completely fills the stroke core as can be seen on the serial sections (**Fig. 1 H-K**).

**Figure 1.**
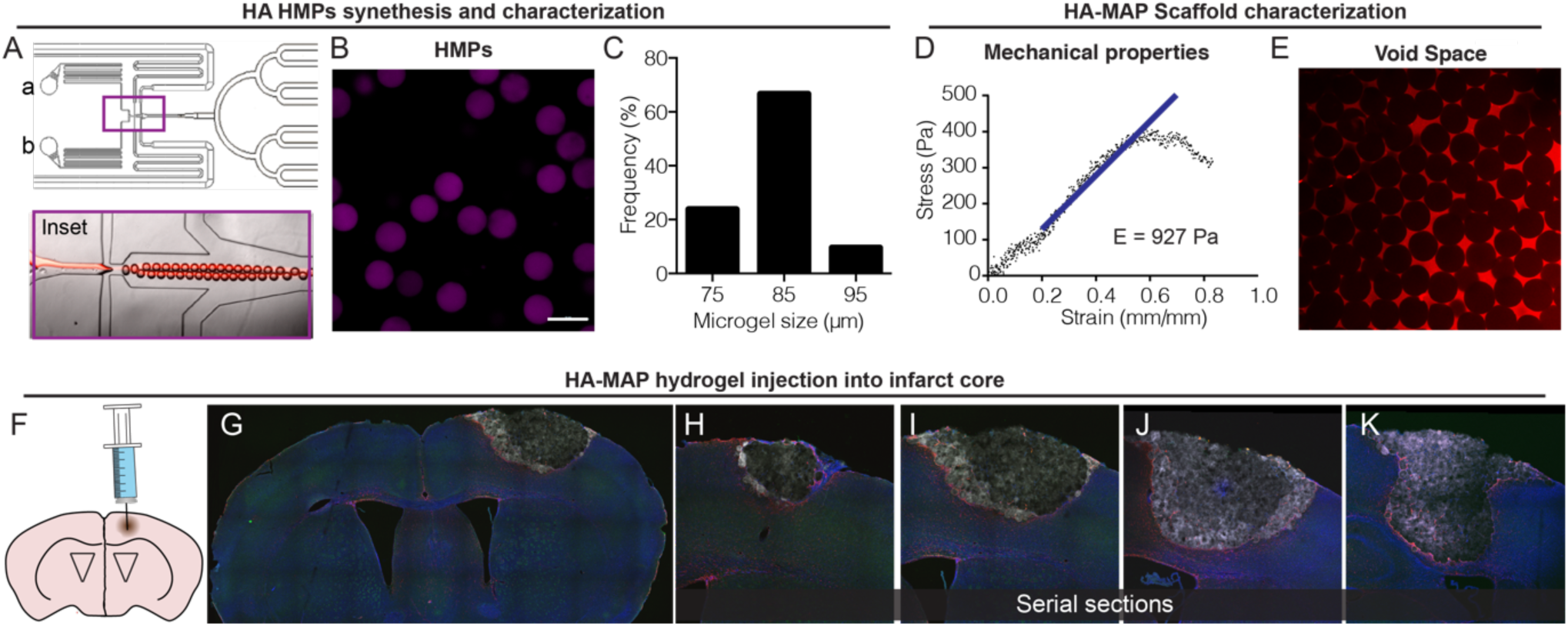
A. Droplet generating flow-focusing microfluidic device used to produce HMPs via a water-in-oil emulsion. B. Image of HMPs in aqueous solution allowed to fully swell after gelation and purification from oil-phase. Scale bar = 100µm C. Histogram of HMP sizes showing majority of HMPs are between 80-90µm. D. Instron mechanical compression testing of MAP scaffold after annealing of HMPs show MAP scaffold exhibits bulk mechanical properties. E. Image of high molecular weight fluorescent dextran between the MAP void spaces showing interconnected porosity. F. Injection schematic showing direct injection into the stroke cavity after PT stroke. G. Full section image of stroked brain treated with MAP showing MAP fills entire stroke cavity.. H-K. Serial section images of single stroked brain treated with MAP with each progressive section ∼300µm apart.

To model ischemic stroke, we used a photothrombotic (PT) stroke model. This model allows examination of the brain tissue’s response at short term (5-15 days post stroke) and long-term time points (30, 120). Time course analysis of macrophage/microglia (IBA1+ cells) and astrocytes (GFAP+ cells) in the infarct and peri-infarct spaces reveals an early peak (7-days) of IBA1+ cells in the core of the infarct, which subsequently subsides reaching a significantly lower level by 30-days (**Sup. Fig. 1A**). The peri-infarct percent area of microglia/macrophages cells and astrocytes increase by 7-days and subsequently plateau through 30-days (**Sup. Fig 1B)**. These data mirror the expected dogma of an early inflammatory response that subsides over time. Analyses of vessels and axons in the peri-infarct space reveal that the vessel area remains relatively constant over time, while the axonal area significantly decreases over time reaching a plateau at 7-days post stroke (**Sup Fig 1C)**. These data show that in the absence of any treatment, little recovery of the peri-infarct axons and vessels is found in this model.

### Injection of MAP hydrogel preserves long term structural integrity and function

In order to understand the feasibility of using HA-MAP hydrogel for brain repair, we first investigated how long the gel lasts in the brain after implantation and if they are any detrimental effects of hydrogel injection over time (**Fig. 2A)**. We injected HA-MAP hydrogel 5-days after stroke and assessed for hydrogel degradation by monitoring hydrogel fluorescence at days 2, 10, 30, 120 days post injection (**Fig. 2B**). We find that HA-MAP hydrogel does not degrade significantly until 120-days post implantation with a significantly decreased fluorescence indicating that the HA-MAP scaffolds can provide long lasting mechanical support to the tissue.

**Figure 2.**
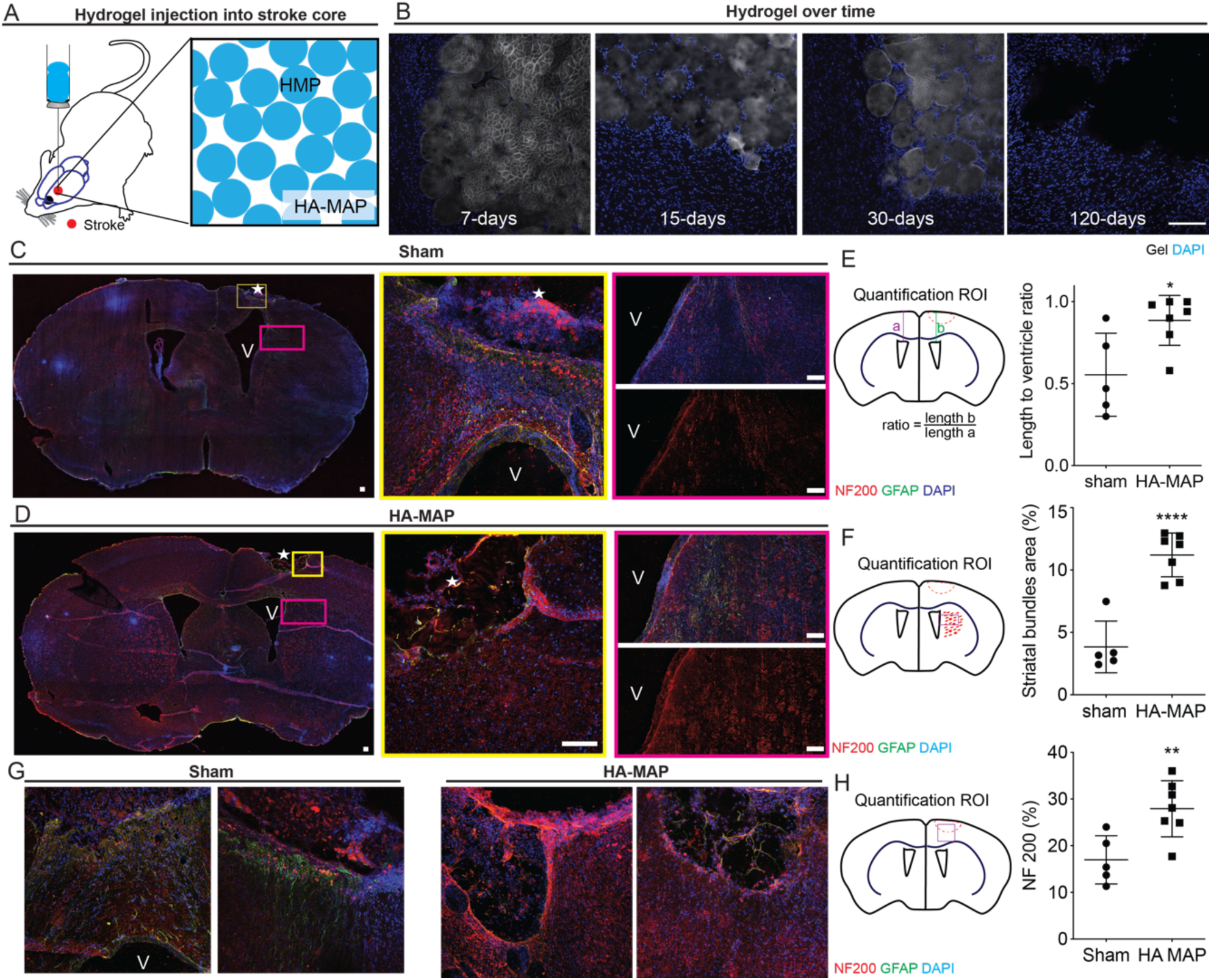
A. Schematic of MAP scaffold directly injected in stroke cavity showing imperfect stacking of HMPs. B. Hydrogel degradation *in vivo* over time after stroke (7, 15, 30, 120 days post stroke). C. IHC images showing cerebral atrophy and reduction of nigrostriatal bundles in the sham condition. V signifies location of ventricle D. IHC images showing preservation of peri-infarct area with minimal cerebral atrophy and nigrostriatal bundles in MAP condition. E. Quantification of cerebral atrophy comparing sham to HA-MAP treated brains. F. Quantification of nigrostriatal bundle area comparing sham to HA-MAP treated brains. G. IHC images comparing the peri-infarct and infarct space of sham to HA-MAP treated brains. H. Quantification of axon percent area above ventricle comparing sham to HA-MAP treated brains. All scale bars = 100µm. Statistical analysis was done in GraphPad Prism. Data were analyzed using a T-test. * indicate P<0.05.

Further analyzing the brain tissue at our longest time point of 120-days post injection, we find that the long-lasting mechanical support is accompanied by maintenance of brain shape over time, which is significantly different from sham (**Fig. 2C & 2D**). In particular, we observe that in the sham group the scar tissue imposes considerable fibrotic response to the brain tissue resulting in cerebral atrophy, whereby the ventricle is pulled upward toward the cortex effectively shrinking the cortex (**Fig. 2E**). In contrast, mice treated with HA-MAP hydrogel 5-days post stroke effectively decreased this response, which significantly prevented the deformation of the stroke cavity. In human patients’ cerebral atrophy in the motor cortex accounts for at least a portion of the worsening motor deficit in stroke patients over time ^8^. We next tested if the density of axons at 120-days post stroke was different between sham and HA-MAP treated groups. We looked at the area between the top of the ventricle to the surface of the cortex. We find that the axonal area (NF200+) was significantly higher for the HA-MAP treated mice compared to sham, further supporting our findings of reduced cerebral atrophy, which is associated with decreased neuronal cell area^9^.

Atrophy is commonly observed in stroke patients and is known to cause effects at regions far away from the stroke core ^9^. Thus, we wanted to get a sense if there are other visible effects of a long-lasting hydrogel in the brain post stroke and reduced cerebral atrophy in mice. We find that the striatal white matter bundles, which include the nigrostriatal bundles^36^, are visible with NF200 staining, and are more preserved in the brains treated with HA-MAP hydrogel compared to sham brains (**Fig. 2F)**. Quantification of the percent positive area of NF200 stain in the bundle area revealed that brains treated with HA-MAP hydrogel more than double the percent area of the sham condition. Nigrostriatal bundles are a dopaminergic pathway that travels from the substantia nigra to the striatum and loss of these bundles is associated with decreased motor function and is commonly observed in Parkinson’s patients ^37, 38^. Parkinson patients also observe significant cerebral atrophy, associated with worsening disease state ^39^. Further, quantification of the percent positive area of NF200 stain in the area directly above the ventricle shows a significant increase in axon area in the HA-MAP treated brains compared to the sham (**Fig 2G, 2H)**. Taken together the reduced cerebral atrophy and thus, preservation of these bundles suggests that the MAP gel would better preserve the motor function associated with these bundles.

### Highly reactive, scar forming, astrocytes are reduced after HA-MAP injection

We next moved to explore the use of HA-MAP hydrogels to modulate astrocyte phenotype. HA-MAP hydrogel was injected 5-days post stroke and tissue was collected 2-days post injection to assess the early reaction to the material (**Fig. 3A,B)**. In the peri-infarct area, we observe a significantly higher percentage of reactive astrocytes in the sham condition expressing pERK1/2 (∼56% compared to ∼25% in MAP) **(Fig. 3C)**, an astrocytic downstream pathway for high reactivity. Moreover, significantly fewer infiltrating astrocytes express pERK1/2 compared to those in the peri-infarct (∼12% in the infarct compared to ∼25% in the peri-infarct) **(Fig. 3D)**. This data suggests that the astrocytes that are infiltrating the MAP gel are less reactive and potentially more pro-reparative. We also observe a significantly higher percentage of reactive astrocytes expressing S100β in the sham condition(∼69%) compared to the 3.5% MAP gel (∼26%), with high expression of S100β considered pathological^40, 41^ **(Fig. 3E)**. Similar to pERK expression, the infiltrating astrocytes expressing S100β (∼15%) were significantly less that in the peri-infarct area (∼26%). Additionally, we used *in situ* hybridization to dive deeper into the reactive astrocyte phenotype (**Fig. 3F)**. Using RNA markers, we probed for C3, a marker shown to be upregulated in neurotoxic reactive astrocytes, and SLC1A2, a marker for all astrocytes ^13^. Using a ratio of C3/SLC1A2 to analyze what percentage of astrocytes are highly reactive, we observed 54.2% of all astrocytes are reactive in the sham condition, while only 25.2% of all astrocytes are reactive in the 3.5% MAP gel peri-infarct condition. Comparing the peri-infarct of the MAP gel (∼25.2%) to the infarct of the MAP gel (∼10.7) once again further shows that the infiltrating astrocytes are less reactive and suggests a pro-regenerative phenotype (**Fig. 3G)**. All these results indicate the MAP gel is able to attenuate astrocyte reactivity just two days after injection by reducing the number of highly reactive neurotoxic astrocytes by greater than 2-fold, promoting a less pro-inflammatory environment in the peri-infarct area and stimulating pro-recovery astrocyte infiltration in the infarct. Since inflammatory responses occur early after injury and initial inflammatory responses have long lasting effects in the tissue ^4^, we expect that early modulation of the astrogliotic response will have long lasting effects.

**Figure 3.**
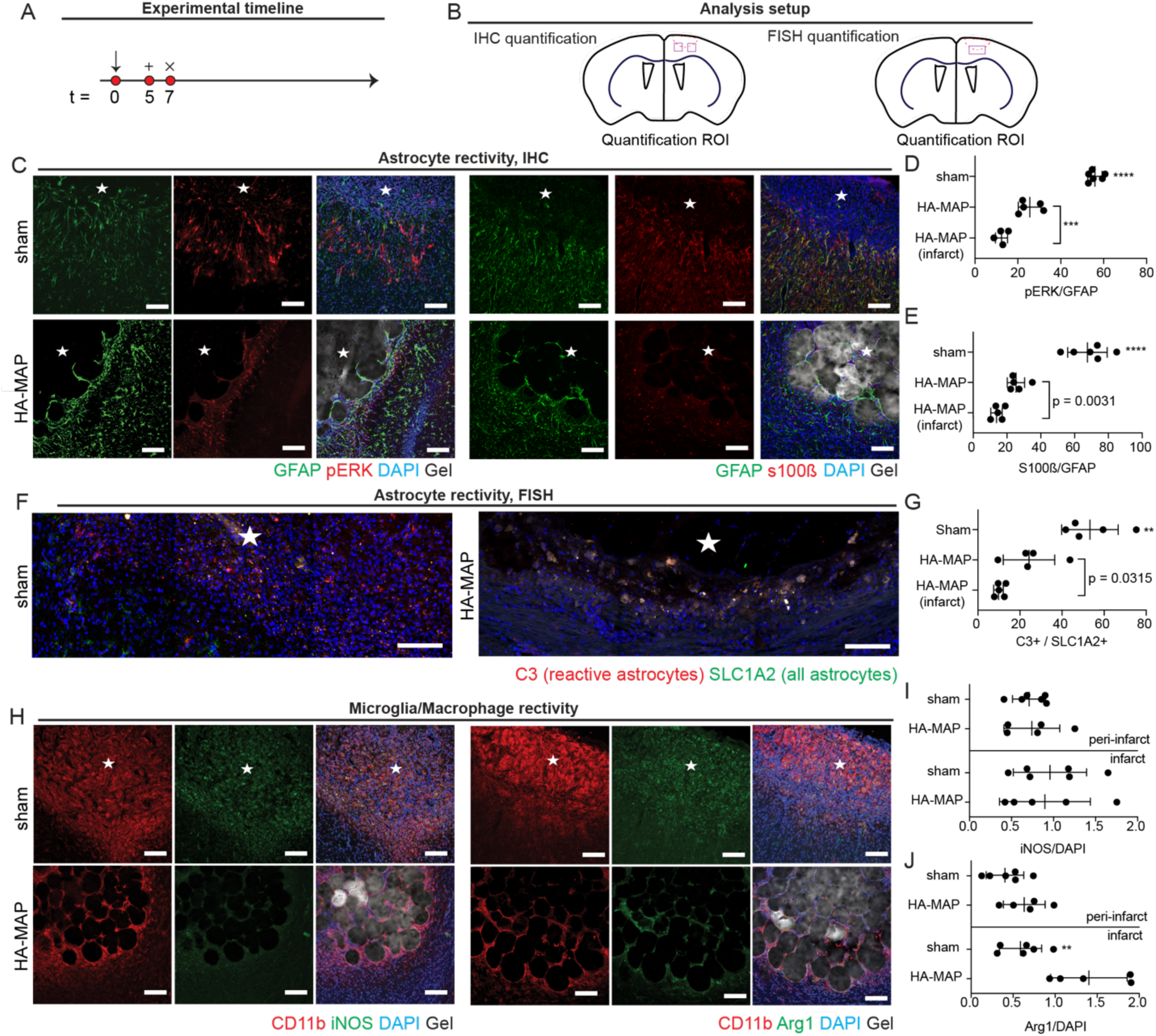
A. Schematic of experimental timeline where arrow signifies stroke day, + signifies injection day, and x signifies sacrifice and analysis time point. B. Schematic of analysis setup showing where sections were imaged and analyzed. C. IHC images showing astrocyte reactivity through pERK, S100β, GFAP. D. Quantification of pERK/GFAP percent of astrocytes that are highly reactive. E. Quantification of S100β/GFAP percent of astrocytes that are highly reactive. F. In situ hybridization images of astrocyte reactivity through C3 and SLC1A2 probing. G. Quantification of C3/SLC1A2 percent of astrocytes that are highly reactive. H. IHC images of microglia reactivity through CD11b, Arg1, and iNOS staining. I. Quantification of iNOS/Dapi for determination of microglial pro-inflammatory phenotype. J. Quantification of Arg1/Dapi for determination of microglial pro-repair phenotype. All scale bar = 100µm. Statistical analysis was done in GraphPad Prism. * indicated data was analyzed using Tukey *post-hoc* test and a 95% confidence interval with P<0.05. Individual p-values indicated T-test.

### Microglia up-regulate pro-repair marker Arg1 in HA-MAP

Microglial polarization directly affects astrocyte reactivity, with microglia that exhibit the more reactive M1 phenotype influencing astrocytes to a more highly reactive state ^13^. Similar to the astrocyte experiments, HA-MAP hydrogel was injected 5-days post stroke and tissue analyzed 2-days later (**Fig. 3H)**. We broadly defined pro-repair microglia/macrophages as expressing Arg1, while pro-inflammatory microglia/macrophages expressing iNOS phenotype ^42^. Interestingly, the percentage of microglia/macrophages expressing the M1 pro-inflammatory phenotype is similar across both conditions in the peri-infarct and in the infarct **(Fig. 3I)**. Further, the percentage of microglia/macrophages expressing the M2 pro-repair phenotype is similar across both conditions in the peri-infarct area. However, the percentage of pro-repair microglia/macrophages in the infarct, where the cells are encapsulated within HA-MAP during injection, is almost 3-fold higher in the MAP gel condition compared to sham condition **(Fig. 3J)**. This suggests that the infarct in the MAP gel has a more pro-reparative environment than the infarct in the sham condition. Confinement of macrophages has been shown to prevent LPS polarized M1 macrophages to activate late stage inflammatory genes ^43^. These findings combined with our findings of increased percentage of Arg1+ cells, suggests that microporous hydrogels formed using ∼80 µm HMPs have pore sizes that can spatially confine the microglia and lower M1 polarization. Chondroitin sulfate proteoglycans (CSPGs) have been linked to decreased regenerative potential in the CNS ^28, 44, 45^. We find that the amount of CSPGs in the HA-MAP treated mice significantly decreases in both the infarct (∼5.3%) and peri-infarct (∼9.8%) compared to sham infarct (∼59.7%) and peri-infarct (∼38%) (**Sup. Fig. 2)**. Several studies have delivered Chondroitinase ABC to digest the high release of CSPG after brain injury and shown behavioral functional benefits^44, 45^. Thus, a material such as HA-MAP that can decrease the CSPG level without additional enzyme delivery would be advantageous.

### Astrocytes continue to infiltrate lesion over time

One key difference between injection of HA-MAP into the stroke cavity compared to any other hydrogel that we have tested ^25, 46-49^ is that it elicits astrocyte infiltration into the stroke cavity. Thus, rather than astrocytes forming a scar around the stroke core, as occurs in sham conditions, astrocytes change their morphology and begin to infiltrate. We observed this previously using a Middle Cerebral Artery occlusion (MCAo) model ^24^ and in data presented here with a PT stroke model, indicating the finding is not model or cortex location specific. To investigate if this change in morphology affects the scar thickness and if astrocyte infiltration is continuous over time, we injected HA-MAP 5-days after stroke and assessed scar thickness and astrocyte infiltration over time (7, 15, 30-days post stroke) by quantifying the thickness of the GFAP+ dense layer surrounding the stroke and infiltration distance of GFAP+ cells starting from the stroke border **(Fig. 4A,B)**. As expected, sham animals have scar thicknesses and peri-infarct % GFAP+ area that increases over time (7-30 days after stroke) (**Fig. 4C**), indicative of a developing scar ^50^. In contrast, injection of HA-MAP hydrogel into the stroke cavity significantly reduces the scar thickness and peri-infarct % GFAP+ area within 2-days of implantation and remains low throughout (**Fig. 4D-F**).

**Figure 4.**
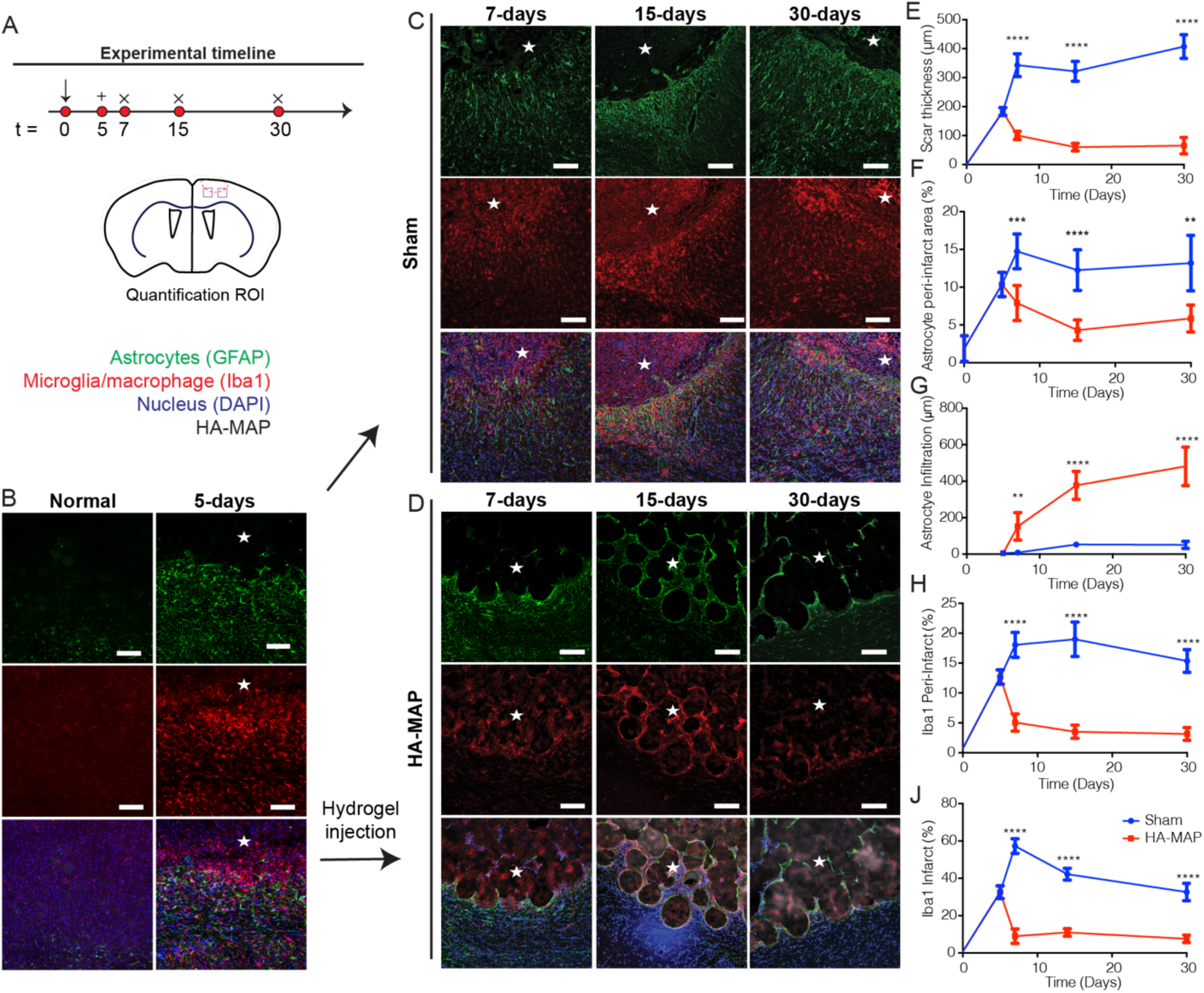
A. IHC images showing reactive astrocytes and microglia of healthy tissue and 5 days post stroke. B. Schematic of experimental timeline where arrow indicates day of stroke, + indicated day of injection, and x indicates days of sacrifice and analysis. C. IHC images of sham stroke tissue showing reactive astrocytes and reactive microglia at 7, 15, and 30 days post stroke through GFAP and IBA-1 staining. D. IHC images of MAP gel treated stroke tissue showing reactive astrocytes and reactive microglia at 7,15, and 30 days post stroke through GFAP and IBA-1 staining. E. Quantification of scar thickness over time through GFAP staining comparing sham and HA-MAP treated brains. F. Quantification of percent area of reactive astrocytes in the peri-infarct through GFAP staining comparing sham and HA-MAP treated brains. G. Quantification of astrocyte infiltration into infarct through GFAP staining comparing sham and HA-MAP treated brains. H. Quantification of percent area of reactive microglia in peri-infarct area through IBA-1 staining comparing sham and HA-MAP treated brains. I. Quantification of percent area of reactive microglia in infarct through IBA-1 staining comparing sham and HA-MAP treated brains. All scale bars = 100µm. Statistical analysis was done in GraphPad Prism. Data were analyzed using a -test and P<0.05.

We next examined astrocyte (GFAP+) cell infiltration over time. We find that astrocyte infiltration begins within 2-days of hydrogel injection and continues to increase from 7-30-days post stroke, reaching close to 500µm into the lesion by day 30 (**Fig. 4G)**. The infiltration pathway closely mimics what we expect is the void structure of the hydrogel (**Fig. 1E)**, indicating that the cells infiltrate through the void space rather than through the HMPs. This infiltration pattern further suggests that significant gel degradation is not required for astrocyte infiltration and that any degraded extracellular matrix (ECM) or new ECM deposition within the void space of HA-MAP does not prevent infiltration of astrocytes. We believe that promoting infiltration of pro-regenerative astrocytes into the infarct can be beneficial towards recovery ^51^. This data also shows that the MAP gel sustains astrocyte infiltration over time without the need for biologic delivery such as growth factors or small molecules.

### Microglia percent decreases over time for hydrogel injection

Next, we examined microglia/macrophage reactivity over time using IBA-1 staining (**Fig. 4D)**. At two days post injection we find a 3-fold or 6-fold decrease in microglia stain for the peri-infarct (**Fig. 4H)** and infarct (**Fig. 4I)** area respectively when comparing sham to HA-MAP treated strokes. As previously mentioned, reactive microglia have been shown to contribute to highly reactive astrocytes ^13^. Further examination into microglial progression shows a very elevated total number of reactive microglia even 30 days post stroke (**Fig. 4H,I)**. Remarkably, the total number of reactive microglia at 30 days post stroke in the infarct of the sham condition is still 3-fold higher than the infarct of the HA-MAP condition at 7 days post stroke, the peak of inflammation.

### Astrocytes and axons co-infiltrate the stroke core after HA-MAP injection

Given the significant impact that HA-MAP has on the phenotype of astrocytes and the number of microglia, we wanted to assess if changes in the stroke environment were accompanied with increases in axonogenesis. As before, HA-MAP hydrogel was injected into the stroke cavity 5-days after stroke and axonal area (NF200+) in the peri-infarct as well as infiltration distance in the infarct quantified (**Fig. 5A,B)**. The peri-infarct percent NF200+ area decreases from 53.2% to 20.4% comparing un-injured brain and stroke brain at 5-days, indicating the rapid loss of neurofilaments in the peri-infarct space after stroke (**Fig. 5C)**. The stroke cavity is devoid of neurofilaments at this timepoint. We find that axons (NF200+) increase in the peri-infarct space following injection of HA-MAP hydrogel, with a significant increase at 7-days post stroke compared with sham (**Fig. 5D-F)**. However, this increase becomes non-significant at the 15- and 30-day timepoints. This data suggests that the lower number of reactive astrocytes combined with the decreased number in macrophage/microglia in the HA-MAP condition, promotes a pro-reparative environment early after HA-MAP injection that promotes axonal sprouting, but this environment cannot be maintained. However, our earlier data at 120-days shows that as cerebral atrophy occurs, axons are maintained in the HA-MAP condition but reduced in sham (**Fig. 2C-H**).

**Figure 5.**
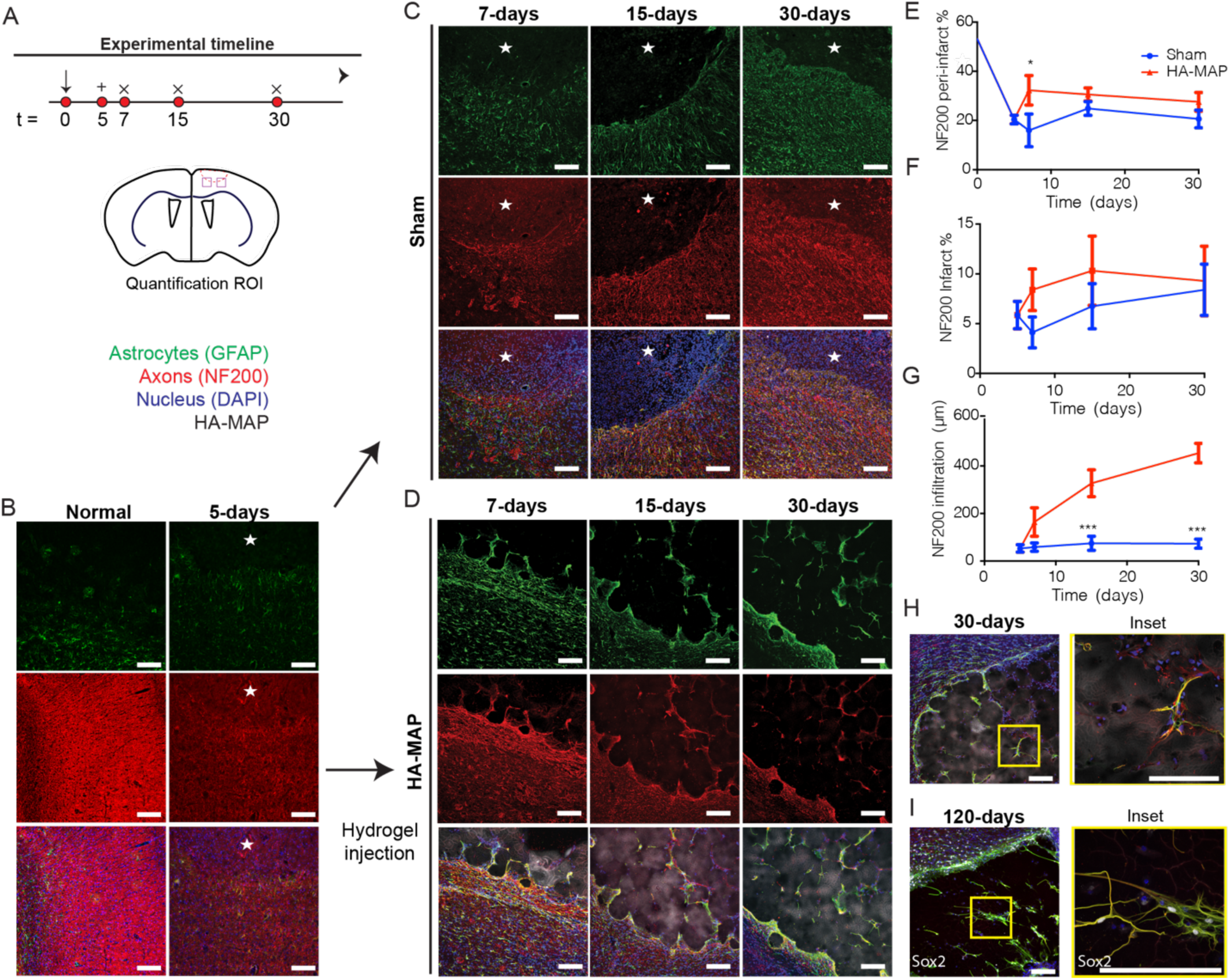
A. IHC images showing reactive astrocytes and axons of healthy tissue and 5 days post stroke through GFAP and NF200 staining. B. Schematic of experimental timeline where arrow indicated day of stroke, + indicated day of injection, and x indicates days of sacrifice and analysis. C. IHC images of sham stroke tissue showing reactive astrocytes and axons at 7, 15, and 30 days post stroke through GFAP and NF200 staining. D. IHC images of MAP gel treated stroke tissue showing reactive astrocytes and axons at 7,15, and 30 days post stroke through GFAP and NF200 staining. E. Quantification of percent area of axons in the peri-infarct through NF200 staining comparing sham and HA MAP treated conditions. F. Quantification of percent area of axons in the infarct through NF200 staining comparing sham and HA MAP treated conditions. G. Quantification of axon infiltration distance into infarct through NF200 staining comparing sham and HA MAP treated conditions. H. IHC images of co-residing astrocytes and axons at 30 days post stroke through GFAP and NF200. I. IHC images of co-residing astrocytes and axons at 120 days post stroke through GFAP and NF200. All scale bars = 100µm. Statistical analysis was done in GraphPad Prism. Data were analyzed using a T-test and P<0.05.

In the infarct cavity we observe axonal infiltration for the HA-MAP hydrogel treated groups, but not in the sham groups. The path of infiltration is similar to what the astrocytes follow, navigating between the HMPs rather than through the HMPs (**Fig. 5G**). The infiltration distance of axons steadily increases over time reaching ∼450µm by day 30 (**Fig. 5G**). Staining for both astrocytes (GFAP) and axons (NF200) revealed that not only do the axons follow the same path, but the axons appear to localize closely with the infiltrating astrocytes (**Fig. 5H & 5I)**. High magnification image analysis revealed that although GFAP stain exists without axonal stain, the reverse is not true, whenever an axonal stain was present it was closely associated with GFAP staining. Examining the HA-MAP hydrogel condition at 4-month reveals that the close association of axons and NF200 is maintained and that it is also accompanied by progenitor cells (SOX2+) (**Fig. 5I)**. Although these images cannot determine the ratio of GFAP+, NF200+, and GFAP+/SOX2+ cells, we likely have all three cell types present within HA-MAP hydrogel at 4 months post injection. Both recruitment of endogenous NPCs and transplantation of exogenous NPCs have been used as strategies to promote stroke repair and shown increase behavioral response^52^. This astrocyte/axon correlation suggests that the astrocytes are crucial for the axon penetration and maintenance, supporting our rationale that promotion of pro-repair astrocyte infiltration will be beneficial for downstream tissue repair. As prior studies have shown, these penetrating astrocytes, in very near proximity to axons could be crucial in forming synapses ^15, 16, 19^ and neuronal circuits directly in the lesion site ^18, 20^, where such recovery is not normally observed after stroke ^4^. Although other molecules and cell types maybe needed to generate the desired circuitry, having astrocytes, axons and NPCs in this same space can be now further manipulated to achieve this. Our results demonstrate that injection of HA-MAP hydrogel into the stroke cavity is generating a permissible environment in the peri-infarct and infarct spaces that leads to axonal infiltration into the stroke core.

### Vessel infiltration does not coincide with astrocyte/axonal infiltration

Similar to axonal density, we observed an initial increase in vessel density (Glut-1+) in the peri-infarct space, which was statistically significant at the 15-day time-point compared to sham mice (**Supplemental Fig. 3A-C**). This result is consistent with other reports of stroke induced angiogenesis in the peri-infarct space ^53^. However, by 30-days post stroke, there is no statistical significance in vessel density between HA-MAP hydrogel injection and sham. Thus, similar to our axonal sprouting data, the increase in angiogenesis suggest that the lower microphage/microglia number and reduced reactive astrocytes in the peri-infarct space generates an environment that further promotes angiogenesis early after HA-MAP gel injection, but that cannot be maintained long term.

After observing significant astrocyte/axonal infiltration into the stroke core, we tested if vessels also infiltrate the stroke core and if they follow a similar infiltration pattern (**Supplemental Fig. 3D**). We find that vessels infiltrate the stroke core, reaching a statistically significant difference of infiltrating vessels between brains treated with HA-MAP and sham groups at 30-days post injection. The infiltration path is similar to that of astrocytes and axons, following the void space between HMPs, but there appears to be no correlation between astrocyte infiltration and vessel infiltration, unlike what was observed for axonal infiltration. The vessel stain Glut-1 does not coincide with the astrocyte stain GFAP, indicating that vessels invade the stroke core independently. Further, we find that the vessel infiltration distance is significantly lower compared to what was found for astrocytes and axons **(Supplemental Fig. 4).** The average infiltration distance at 30-days for vessels is ∼207µm while it is ∼467µm for axons and astrocytes. This difference in infiltration distance is perplexing as HA-MAP hydrogel contains no releasable bioactive signals that would promote the infiltration of astrocytes and axons but not vessels. A few possibilities for the reduced infiltration distance for vessels are the physical properties of the gel, mechanical properties, and porosity. It is difficult to know what the mechanical properties of the gel are over time; however, the initial HA-MAP gel Young’s modulus is ∼1000Pa. It is possible that the mechanical modulus of the gel is not optimal for vessel infiltration, though others, including our lab, have used soft hydrogels for vascularization in the brain ^30^, skin ^30^, and bone ^54^. Thus, mechanical properties are not likely to be the reason. The porosity of the scaffold is another possibility; HA-MAP gels with HMPs of 80-100µm in diameter have a median pore area of ∼200µm^2^ which would correspond to 15-20µm when measured per z-stack ^30, 55^. Although we have previously shown that PEG-MAP gels with 80µm diameter HMPs promote vascularization and rapid wound closure in skin wounds ^30^, others have shown that in cardiac engineering there is limited vessel penetration in scaffolds with 20µm pores^56^. Thus, the effect in pore size in vascularization maybe tissue specific. Regardless of the reason, we have previously demonstrated that a robust vascular network within the stroke, promotes the formation of a neurovascular niche that leads to effective axonogenesis and behavioral improvement ^25^. Thus, efforts should be made to introduce pro-angiogenic cues into HA-MAP to promote vascularization.

An additional rationale for further promoting vascularization is the fact that astrocytes and axons stopped infiltrating 200µm away from vessels, which is the oxygen diffusion limit ^57^. Thus, lack of vascular infiltration, likely limited the range of astrocyte and axonal infiltration.

### Moderate hydrogel stiffness changes do not affect astrocyte behavior

Biomaterial and substrate stiffness has been previously shown to regulate astrocyte reactivity with softer substrates promoting astrocyte quiescence, suggesting that softer substrates should be used to better modulate astrocytes following stroke^58^. Stiffness has also been implicated in axonal sprouting, vessel sprouting and microglia polarization. We tested a moderately stiffer HA-MAP hydrogel (3.5% = 1000Pa, 4.5% = 1500Pa) to determine if this stiffness change can affect astrocyte behavior. All other parameters, RGD concentration, HMP diameter, and void fraction were kept constant. Overall, the 4.5% MAP hydrogel produced very similar results compared to the 3.5% hydrogel for astrocyte infiltration and scar thickness (**S. Fig. 5A & 5B)**. However, we did observe an increased number of reactive microglia in the infarct and in the peri-infarct at 15 days post stroke, but no significant differences were observed at 30 days post stroke **(S. Fig. 5C & 5D)**. Interestingly, these increased microglia response may have caused some downstream effects as we observe a slight decrease in axon penetration compared to the 3.5% condition (S. **Fig. 5E)**. It is possible that the increased number of microglia in the 4.5% condition created a more cytotoxic environment that affected the axon penetration. Overall, the difference between 1000Pa and 1500Pa modulus gels may be too small to see more significant responses.

### Porosity, hyaluronic acid, and RGD are crucial for astrocyte infiltration

We next wanted to examine if the observed responses are specific to the HA-MAP formulation tested. HA-MAP are produced from ∼80µm HMPs, which contain HA (70,000 Da, 3.5%), RGD (500µM), K and Q peptides (250µM), and 7.8mM of MMP crosslinker. The MAP gel is crosslinked using FXIIIa (5U/mL) and 1U/mL of thrombin. To test the role of microstructure on the observed findings, we compared the results from HA-MAP to those of a hydrogel with identical composition but crosslinked as a bulk gel (non-porous) (**Fig. 6A,B)**. As a comparison we tested the ability of these gels to promote astrocyte infiltration, reduce scar thickness and reduce the number of microphage/microglia in the peri-infarct and infarct spaces. We find that the scar thickness in the non-porous group is significantly reduced compared to sham but significantly larger compared to HA-MAP **(Fig. 6B**). There is also no infiltration of astrocytes into the stroke cavity, suggesting that the microstructure of MAP is critical to this process. Microglia/macrophage area was significantly decreased in the infarct but not the peri-infarct compared to sham; however, the decrease was not as great as that observed with HA-MAP.

**Figure 6:**
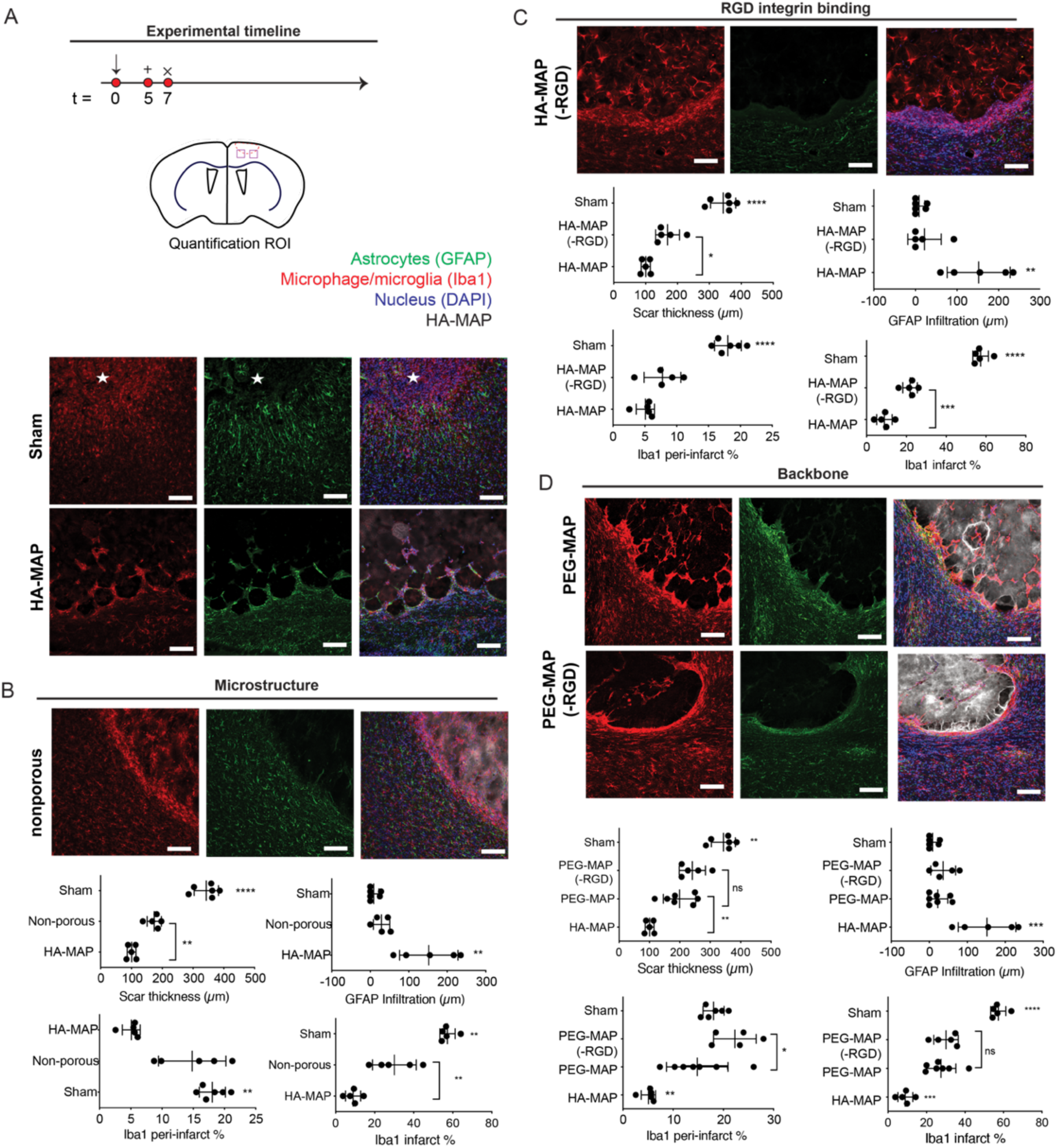
A. Schematic of experimental timeline where arrow indicates day of stroke, + indicates day of injections, and x indicates day of sacrifice and analysis. IHC images for sham and HA MAP treated brains at 7 days post stroke staining for GFAP and IBA-1. B. IHC images of non-porous treated brains staining for GFAP and IBA-1. Quantification of reactive astrocytes and reactive microglia testing for scar thickness, astrocyte infiltration distance, IBA-1 peri-infarct percent area, and IBA-1 infarct percent area to analyze the role of hydrogel microstructure through GFAP and IBA-1 staining. C. IHC images of HA MAP (-RGD) treated brains staining for GFAP and IBA-1. Quantification of reactive astrocytes and reactive microglia testing for scar thickness, astrocyte infiltration distance, IBA-1 peri-infarct percent area, and IBA-1 infarct percent area to analyze the role of RGD integrin binding through GFAP and IBA-1 staining. D. IHC images of PEG-MAP and PEG-MAP (-RGD) treated brains staining for GFAP and IBA-1. Quantification of reactive astrocytes and reactive microglia testing for scar thickness, astrocyte infiltration distance, IBA-1 peri-infarct percent area, and IBA-1 infarct percent area to analyze the role of hydrogel backbone through GFAP and IBA-1 staining. All scale bars = 100µm. Statistical analysis was done in GraphPad Prism. Data were analyzed using a one way analysis of variance followed by a Tukey *post-hoc* test and a 95% confidence interval. * indicate P<0.05.

We next tested the effect of the integrin binding ligand RGD. Injection of HA-MAP(-RGD) revealed that the scar thickness was again lower than sham, but not as low as HA-MAP. Similar to HA non-porous there was no infiltration of astrocytes into the stroke cavity, pointing to the importance of RGD in this process. The number of reactive microglia for HA-MAP(-RGD) is similar to HA-MAP for the peri-infarct space, but significantly higher for infarct (**Fig. 6C**). Overall, we see that RGD has a major effect in the ability of HA-MAP to modulate astrocyte and microglia phenotype.

Last, we tested the role of the hydrogel backbone and compared HA to polyethylene glycol (PEG). PEG-vinyl sulfone (20,000Da) was used to generate HMPs using the same concentration of K, Q and MMP peptides. PEG HMPs were generated with RGD or no RGD. PEG HMPs have the same diameter as HA HMPs (**Supplemental Fig. 6)** and the storage modulus is the same as HA-MAP (**Supplemental Fig. 6**). We find that having a PEG backbone in the HMPs significantly reduces the effect observed with HA-MAP. Scar thickness is significantly thinner than sham, but larger than HA-MAP and there is no significant difference between PEG-MAP and PEG-MAP(-RGD) (**Fig. 6D)**. Astrocyte infiltration in the PEG-MAP or PEG-MAP(-RGD) conditions is also close to zero. Finally, the microphage/microglia area is significantly decreased in the infarct for both PEG-MAP or PEG-MAP(-RGD), but not statistically different from sham in the peri-infarct space. Overall, we find that HA is an essential component to promote infiltration of astrocytes into the infarct core.

Taken together this data shows that HA-MAP composition and microstructure is essential for the observed astrocyte infiltration, reduced scar thickness, and reduced microphage/microglia in the peri-infarct and infarct spaces. No other condition tested was equally effective as HA-MAP. Importantly, all the conditions that used HA as the backbone significantly decreased the scar thickness compared to conditions that used PEG as the backbone, demonstrating that the biological activity of HA plays an important role in this process. After stroke, CD44 expression is elevated in reactive astrocytes ^59^. Thus, receptor mediated astrocyte/HA binding maybe responsible for the infiltration in HA-MAP but not PEG-MAP. Previous studies implanting porous HA-RGD hydrogels into cortex, demonstrated that RGD increased astrocyte infiltration into the cortex wound ^60^. These results agree with our finding that HA-MAP but not HA-MAP(-RGD) promoted astrocyte infiltration. It is surprising that PEG-MAP and HA-MAP resulted in different tissue responses. These materials have the same microstructures and same concentration of RGD. Given the large number of studies conducted using PEG hydrogels for tissue repair applications, it is possible that the results observed here are particular to brain and not generalizable to all tissues. In our own laboratory, we have used PEG-MAP to treat skin wounds and shown substantial cellular infiltration and wound closure just 5-days after implantation ^30^.

## Conclusions

Astrocytes have previously been thought to exacerbate the inflammation after stroke and be detrimental to recovery ^5^, however, recent studies show astrocytes with pro-regenerative phenotypes can be beneficial to the repair process^13^. When injected directly into the stroke cavity, the HA-MAP hydrogel, a microporous scaffold assembled from the imperfect stacking of uniform particle hydrogels produced in a flow-focusing microfluidic device, drastically affects the phenotype and reactivity state of the astrocytes just two days after injection. Using markers of highly reactive astrocytes, we show that the HA MAP hydrogel significantly reduces highly reactive astrocytic phenotype compared to sham, while also promoting the infiltration of less reactive astrocytes into the lesion. Examination into microglial polarity suggested a more pro-repair environment compared to sham. Taken together, the HA-MAP is able to modulate the brain’s inflammatory response to the stroke by reducing highly reactive astrocytes and promoting infiltration of cells with a pro-reparative phenotype. The HA-MAP hydrogel is able to sustain decreased inflammation over time by maintaining a reduced astrocytic scar, preventing a large influx of reactive microglia/macrophages, while extending astrocyte infiltration into the lesion. The ability of the HA-MAP gel to modulate inflammation could transform treatments of severe inflammation in the brain.

By decreasing inflammation and promoting astrocytic infiltration, the HA MAP hydrogel is able to influence axonal penetration at the injured site. Every axon penetrating in the lesion is observed to be co-residing next to an infiltrating astrocyte. Meanwhile in the sham we observe minimal astrocyte and axon infiltration. This suggests that the less reactive astrocytes infiltrating into the lesion are partaking in the recovery process by supporting axonal infiltration. Thus, the anti-inflammatory effects of the HA-MAP hydrogel has downstream benefits by promoting increasing axonal penetration over time. Looking further downstream, we observe that the injection of the HA-MAP hydrogel is able to provide mechanical support to the surrounding brain tissue by preventing the fibrotic pulling of the ventricle known as cerebral atrophy. With the combination of early on decreased inflammation and increased mechanical support provided by the HA-MAP gel, the striatal pathway bundles, associated with motor function ^37^, are much better preserved compared to sham. This suggests that the ability of the HA-MAP gel to modulate inflammation and support the tissue long term has significant downstream benefits and can lead to improved motor function.

Systematically testing effect of microstructure, RGD, and polymer backbone, we show that all three play crucial roles in modulating astrocyte infiltration. All injected hydrogels decreased inflammation to some extent, however, the HA-MAP hydrogel significantly decreased the reactive microglial and astrocytic population the most. Moreover, only the HA-MAP hydrogel was able to promote astrocyte infiltration into the lesion. This suggests that porosity and RGD are required for astrocyte infiltration, while receptor mediated astrocyte/HA binding could also be critical to infiltration.

The HA MAP hydrogel does have vascular limitations and further engineering is required of the material to improve the vascular response. Vascularization is critical to providing oxygen and nutrients to neighboring cells as its been shown that the oxygen diffusion limit is ∼200µm. We observe the astrocytes and axons to be infiltrating ∼200µm further than the infiltrating vessels. This suggests that the limiting factor to further astrocyte/axonal infiltration is the vessel response. Likely introducing delivery of angiogenic factors is required to improve the vasculature, infiltration and thereby, improving astrocyte/axonal infiltration.

## Material and Methods

### Microgel Production and Purification

Microfluidic devices^30^ and 3.5% microgels^61^ were produced as previously described. Briefly, hyaluronic acid (HA) functionalized with an acrylate was dissolved at 7% (w/v) in 0.3M triethyloamine (TEOA) pH 8.8 and pre-reacted with K-peptide (Ac-FKGGERCG-NH_2_) and Q-peptide (Ac-NQEQVSPLGGERCG-NH_2_) at a final hydrogel concentration of 250µM, and RGD (Ac-RGDSPGERCG-NH_2_) at a final hydrogel concentration of 500μM. Concurrently, the cross-linker solution was prepared by dissolving the di-thiol matrix metalloproteinase (MMP) sensitive peptide (Ac-GCRDGPQGIWGQDRCG-NH_2_, Genscript) in distilled water at 7.8 mM and reacted with 10 μM Alexa-Fluor 647-maleimide (Life-Technologies) for 5 minutes. These solutions were mixed in a flow focusing microfluidic device and then immediately pinched by 1% span-80 in heavy mineral oil to form microspheres. These microspheres were collected and allowed to gel overnight at room temperature to form microgels. The microgels were then purified by repeated washes with HEPES buffer (pH 7.4 containing 10mM CaCl_2_) and centrifugation. 4.5% microgels were produced in the same manner, however, the HA-acrylate precursor solution was dissolved at 9% (w/v). The HA without RGD microgels were produced with the same precursor solution as the 3.5% HA, however, no RGD was added. The PEG microgels were produced as previously described^30^. Briefly, 4-arm PEG-Vinyl sulfone (Jenkem) was dissolved at 10% (w/v) and the peptide concentrations were the same as the HA solutions.

### Generation of scaffold from microgels and mechanical testing

The microgels were pelleted by centrifuging at 18,000 G and the supernatant was discarded to form a concentrated solution of microgels. 5 U/mL of FXIII and 1 U/mL of Thrombin were combined in the presence of 10mM Ca^2+^ with the pelleted µgels and allowed to incubate at 37 °C for 90 minutes between two glass slides (1mm thickness) surface coated with sigmacote (Sigma-Aldrich). The mechanical testing on the hydrogel scaffolds was done using a 5500 series Instron. After annealing, the scaffolds were allowed to swell in HEPES buffer saline for 4 hours at room temperature. A 2.5N load cell with a 3.12mm tip in diameter was used at a compression strain rate of 1mm/min and the hydrogel scaffold was indented 0.8mm or 80% of its total thickness.

### Nanoporous Hydrogel Production

Nanoporous hydrogel precursor solutions were exactly the same as the microgel precursor solutions. Additionally, the same enzyme sensitive di-thiol cross-linker solution was prepared. These two solutions were thoroughly mixed in an Eppendorf tube by vortexing and pipetting. 5 U/mL of FXIII and 1 U/mL of Thrombin were added to the solution and the nanoporous hydrogel was allowed to gel *in situ* via the same Michael type addition in which the microgels were individually formed.

### In vivo Photothrombotic Stroke Model

Animal procedures were performed in accordance with the US National Institutes of Health Animal Protection Guidelines and the University of California Los Angeles Chancellor’s Animal Research Committee. A cortical photothrombotic stroke was induced on 8-12 week male C57BL/6 mice obtained from Jackson laboratories (Bar Harbor, ME). The mice were anesthetized with 2.5% isoflurane and placed onto a stereotactic setup. The mice were kept at 2.5% isoflurane in N2O:O2 for the duration of the surgery. A midline incision was made and Rose Bengal (10 mg/mL, Sigma-Aldrich) was injected intraperitoneally at 10 μL/g of mouse body weight. After 5 minutes of Rose Bengal injection, a 2-mm diameter cold fiberoptic light source was centered at 0 mm anterior/1.5 mm lateral left of the bregma for 18 minutes and a burr hole was drilled through the skull in the same location. All mice were given sulfamethoxazole and trimethoprim oral suspension (TMS, 303 mL TMS/250 mL H20, Amityville, NY) every 5 days for the entire length of the experiment. Five days following stroke surgery, microgels with FXIII were loaded into a Hamilton syringe (Hamilton Reno, NV) connected to a pump and 6 μL of microgels were injected into the stroke cavity using a 30-gauge needle at stereotaxic coordinates 0.26 mm anterior/posterior (AP), 3 mm medial/lateral (ML), and 1 mm dorsal/ventral (DV) with an infusion speed of 1μL/min. The needle was withdrawn from the mouse brain 5 minutes after the injection to allow for microgel annealing. For each condition a minimum of 5 mice was used.

Seven days, Fifteen days, thirty days, and one hundred twenty days following stroke, mice were sacrificed via transcardial perfusion of 0.1 M PBS followed by 40 mL of 4 (w/v) % PFA. The brains were isolated and post-fixed in 4% PFA overnight and submerged in 30 (w/v) % sucrose solution for 24 hours. Tangential cortical sections of 30 μm-thickness were sliced using a cryostat and directly mounted on gelatin-subbed glass slides for immunohistological staining of GFAP (glial fibrillary acidic protein, Abcam, Cambridge, MA, USA) for astrocytes, IBA-1 (ionized calcium binding adaptor molecule, Abcam, Cambridge, MA, USA) for microglial cells, Glut-1 (Glucose Transporter-1, Abcam, Cambridge, MA, USA) for endothelial cells, NF200 (Neurofilament 200, Abcam, Cambridge, MA, USA) for axonal processes, pERK (Cell Signaling) for highly reactive astrocytes, S100β (thermofisher) for highly reactive astrocytes, CD11b (Abcam) for immune cells, Arginase 1 (Santa Cruz) for Pro-repair microglia/macrophages, NOS2 (Santa Cruz) for pro-inflammatory microglia/macrophages, Sox2 (Santa Cruz) for neural progenitor cells, and DAPI (1:500 Invitrogen) for nuclei. Primary antibodies (1:100) were incubated overnight at 4°C and secondary antibodies (1:1000) were incubated at room temperature for two hours. A Nikon C2 confocal microscope was used to take fluorescent images.

### Image analysis

The IBA-1, GFAP, pERK, S100β, CD11b, Arg1, iNOS, NF200, and Glut-1 astrocytic (GFAP) and positive area in the infarct and peri-infarct areas were quantified in 4 to 8 randomly chosen regions of interest (ROI of 0.3 mm^2^) at a maximum distance of 300µm from the infarct for the peri-infarct analysis. In each ROI, the positive area was measured

### Ventricular Hypertrophy and Nigrostriatal Bundles analysis (Figure 2)

Cryofrozen sections were allowed to thaw at room temperature. Sections were washed with PBS for 5 minutes with 3 repetitive washes. Sections were incubated with a 10% donkey serum and PBS with 0.3% triton at room temperature for one hour. The liquid was wicked away and primary antibodies of GFAP (glial fibrillary acidic protein, Abcam, Cambridge, MA, USA) for astrocytes and NF200 (Neurofilament 200, Abcam, Cambridge, MA, USA) for axonal processes at 1:100 dilution in PBS with 0.3% triton and 10% donkey serum were added and incubated overnight at 4°C. The next day the primaries were washed with 3 repeated PBS washes of 5 minutes each. Secondary antibodies with donkey hosts along with Dapi at 1:1000 dilution in PBS with 0.3% triton and 10% donkey serum were added and incubated at room temperature for 2 hours. After two hours, the secondary antibodies were washed away with 3 repeated PBS washes of 5 minutes each. The sections were allowed to dry at room temperature and mounted using DPX mounting medium. Analyses were performed on microscope images of 3 coronal brain levels at +0.80 mm, −0.80 mm and −1.20 mm according to bregma, which consistently contained the cortical infarct area. Large scale 10x images of each section was taken and analyzed for ventricular hypertrophy. The ratio of the ipsilateral length from the top of the section to the top of the ventricle was divided by the ratio of the contralateral length from the top of the section to the top of the ventricle was taken to get a quantitative number for the ventricular hypertrophy. Large scale 20x images were taken by the side of ventricle to analyze for nigrostriatal bundle area. Using pixel threshold on 8-bit converted images using ImageJ (Image J v1.43, Bethesda, Maryland, USA) and expressed as the area fraction of positive signal per area (%). Values were then averaged across all areas and sections, and expressed as the average positive area per animal. The percent area positive for NF200 was analyzed 0-1mm out from the ventricle.

### Astrocyte Reactivity, IHC (figure 3)

Cryofrozen sections were allowed to thaw at room temperature. Sections were washed with PBS for 5 minutes with 3 repetitive washes. Sections were incubated with a 10% donkey serum and PBS with 0.3% triton at room temperature for one hour. The liquid was wicked away and primary antibodies of GFAP (glial fibrillary acidic protein, Abcam, Cambridge, MA, USA) for astrocytes and pERK (Cell Signaling) for highly reactive astrocytes at 1:100 dilution in PBS with 0.3% triton and 10% donkey serum were added and incubated overnight at 4°C. The next day the primaries were washed with 3 repeated PBS washes of 5 minutes each. Secondary antibodies with donkey hosts along with Dapi at 1:1000 dilution in PBS with 0.3% triton and 10% donkey serum were added and incubated at room temperature for 2 hours. After two hours, the secondary antibodies were washed away with 3 repeated PBS washes of 5 minutes each. The sections were allowed to dry at room temperature and mounted using DPX mounting medium. Analyses were performed on microscope images of 3 coronal brain levels at +0.80 mm, −0.80 mm and −1.20 mm according to bregma, which consistently contained the cortical infarct area. Each image represents a maximum intensity projection of 10 to 12 Z-stacks, 1 µm apart, captured at a 20x magnification with a Nikon C2 confocal microscope using the NIS Element software. For the sham sections, using ImageJ and converting to 8-bit, a ratio of positive pERK area divided by positive GFAP area within the same area was taken to get percent of reactive astrocytes that are highly reactive in the peri-infarct area 0-300µm from the infarct border. For the HA MAP sections, the peri-infarct were analyzed similarly to sham. The infarct was analyzed by taking 0-100µm infiltration into the lesion. S100β (thermofisher) was stained, imaged, and analyzed similarly to pERK.

### Astrocyte Reactivity, HCR (figure 3)

In situ hybridization probes for C3 and SLC1A2 were purchased from Molecular Instruments and the protocol published on the molecular instrument website was used to probe the sections. Analyses were performed on microscope images of 3 coronal brain levels at +0.80 mm, −0.80 mm and −1.20 mm according to bregma, which consistently contained the cortical infarct area. Large scale 40x images were taken using a Nikon C2 confocal along the stroke border. The images were than analyzed similar to pERK and S100β.

### Microglial Reactivity (figure 3)

Cryofrozen sections were allowed to thaw at room temperature. Sections were washed with PBS for 5 minutes with 3 repetitive washes. Sections were incubated with a 10% donkey serum and PBS with 0.3% triton at room temperature for one hour. The liquid was wicked away and primary antibodies of CD11b (Abcam) for immune cells, Arginase 1 (Santa Cruz) for Pro-repair microglia/macrophages, NOS2 (Santa Cruz) for pro-inflammatory microglia/macrophages at 1:100 dilution in PBS with 0.3% triton and 10% donkey serum were added and incubated overnight at 4°C. The next day the primaries were washed with 3 repeated PBS washes of 5 minutes each. Secondary antibodies with donkey hosts along with Dapi at 1:1000 dilution in PBS with 0.3% triton and 10% donkey serum were added and incubated at room temperature for 2 hours. After two hours, the secondary antibodies were washed away with 3 repeated PBS washes of 5 minutes each. The sections were allowed to dry at room temperature and mounted using DPX mounting medium. Analyses were performed on microscope images of 3 coronal brain levels at +0.80 mm, −0.80 mm and −1.20 mm according to bregma, which consistently contained the cortical infarct area. Each image represents a maximum intensity projection of 10 to 12 Z-stacks, 1 µm apart, captured at a 20x magnification with a Nikon C2 confocal microscope using the NIS Element software. In the peri-infarct for both sham and HA MAP, the ratio of the positive area of iNOS or Arg1 was divided by the positive area for CD11b from 0-300µm from the infarct border. In the infarct, the ratio of the positive area of iNOS or Arg1 was divided by the positive area for CD11b from 0-300µm from the infarct border.

### Astrocyte and Microglial progession analysis (Figure 4)

Cryofrozen sections were allowed to thaw at room temperature. Sections were washed with PBS for 5 minutes with 3 repetitive washes. Sections were incubated with a 10% donkey serum and PBS with 0.3% triton at room temperature for one hour. The liquid was wicked away and primary antibodies of GFAP (glial fibrillary acidic protein, Abcam, Cambridge, MA, USA) for astrocytes, IBA-1 (ionized calcium binding adaptor molecule, Abcam, Cambridge, MA, USA) for reactive microglia/macrophages at 1:100 dilution in PBS with 0.3% triton and 10% donkey serum were added and incubated overnight at 4°C. The next day the primaries were washed with 3 repeated PBS washes of 5 minutes each. Secondary antibodies with donkey hosts along with Dapi at 1:1000 dilution in PBS with 0.3% triton and 10% donkey serum were added and incubated at room temperature for 2 hours. After two hours, the secondary antibodies were washed away with 3 repeated PBS washes of 5 minutes each. The sections were allowed to dry at room temperature and mounted using DPX mounting medium. Analyses were performed on microscope images of 3 coronal brain levels at +0.80 mm, −0.80 mm and −1.20 mm according to bregma, which consistently contained the cortical infarct area. Each image represents a maximum intensity projection of 10 to 12 Z-stacks, 1 µm apart, captured at a 20x magnification with a Nikon C2 confocal microscope using the NIS Element software. Scar thickness was analyzed as distance from stroke border using astrocyte morphology changes to signify end of scar. Astrocyte infiltration was measured as longest infiltrating astrocyte from stroke border. In the peri-infarct for both sham and HA MAP, the percent positive for IBA-1 was measured 0-300µm from the infarct border. In the infarct, the percent positive for IBA-1 was measured 0-300µm from the infarct border.

### Axon progression (Figure 5)

Cryofrozen sections were allowed to thaw at room temperature. Sections were washed with PBS for 5 minutes with 3 repetitive washes. Sections were incubated with a 10% donkey serum and PBS with 0.3% triton at room temperature for one hour. The liquid was wicked away and primary antibodies of GFAP (glial fibrillary acidic protein, Abcam, Cambridge, MA, USA) for astrocytes and NF200 (Neurofilament 200, Abcam, Cambridge, MA, USA) for axonal processes at 1:100 dilution in PBS with 0.3% triton and 10% donkey serum were added and incubated overnight at 4°C. The next day the primaries were washed with 3 repeated PBS washes of 5 minutes each. Secondary antibodies with donkey hosts along with Dapi at 1:1000 dilution in PBS with 0.3% triton and 10% donkey serum were added and incubated at room temperature for 2 hours. After two hours, the secondary antibodies were washed away with 3 repeated PBS washes of 5 minutes each. The sections were allowed to dry at room temperature and mounted using DPX mounting medium. Analyses were performed on microscope images of 3 coronal brain levels at +0.80 mm, −0.80 mm and −1.20 mm according to bregma, which consistently contained the cortical infarct area. Each image represents a maximum intensity projection of 10 to 12 Z-stacks, 1 µm apart, captured at a 20x magnification with a Nikon C2 confocal microscope using the NIS Element software. Axon infiltration was measured as longest infiltrating neurofilament from stroke border. In the peri-infarct for both sham and HA MAP, the percent positive for NF200 was measured 0-300µm from the infarct border. In the infarct, the percent positive for NF200 was measured 0-300µm from the infarct border.

### Vessel Progression (Supplemental Figure)

Cryofrozen sections were allowed to thaw at room temperature. Sections were washed with PBS for 5 minutes with 3 repetitive washes. Sections were incubated with a 10% donkey serum and PBS with 0.3% triton at room temperature for one hour. The liquid was wicked away and primary antibodies of GFAP (glial fibrillary acidic protein, Abcam, Cambridge, MA, USA) for astrocytes and Glut-1 (Glucose Transporter-1, Abcam, Cambridge, MA, USA) for endothelial cells at 1:100 dilution in PBS with 0.3% triton and 10% donkey serum were added and incubated overnight at 4°C. The next day the primaries were washed with 3 repeated PBS washes of 5 minutes each. Secondary antibodies with donkey hosts along with Dapi at 1:1000 dilution in PBS with 0.3% triton and 10% donkey serum were added and incubated at room temperature for 2 hours. After two hours, the secondary antibodies were washed away with 3 repeated PBS washes of 5 minutes each. The sections were allowed to dry at room temperature and mounted using DPX mounting medium. Analyses were performed on microscope images of 3 coronal brain levels at +0.80 mm, −0.80 mm and −1.20 mm according to bregma, which consistently contained the cortical infarct area. Each image represents a maximum intensity projection of 10 to 12 Z-stacks, 1 µm apart, captured at a 20x magnification with a Nikon C2 confocal microscope using the NIS Element software. Vessel infiltration was measured as longest infiltrating vessel from stroke border. In the peri-infarct for both sham and HA MAP, the percent positive for Glut1 was measured 0-300µm from the infarct border. In the infarct, the percent positive for Glut1 was measured 0-300µm from the infarct border.

### Statistical Analysis

Statistical analyses were performed as previously described^49^. For histology a minimum of n=5 was used. The results are expressed as mean ± s.e.m. A *P* value < 0.05 was considered statistically significant.

## Acknowledgements

The authors would like to acknowledge Dr. Amy Gleichman for her expertise on astrocytes and helpful discussion. The authors would like thank Dr. Irene Llorente for her help on quantification methods. The authors would like to acknowledge Dr. Lina Nih for her assistance with tissue collection and helpful discussions. The authors would like thank Kat Wilson for her help producing HMPs. The authors would like to thank the entire Carmichael lab at UCLA for their stroke expertise and help. We would like to acknowledge the National Institutes of Health and the National Institute for Neurological Diseases and Stroke for funding (R01NS094599).

## Author contributions

E.S., STC, T.S. designed the experiments. E.S., A.Y., and J.C. performed and analyzed data. E.S., STC, T.S. wrote the manuscript with input from all authors.

## Supplementary Material

### Supplementary Figures

**Supplemental Figure 1.**
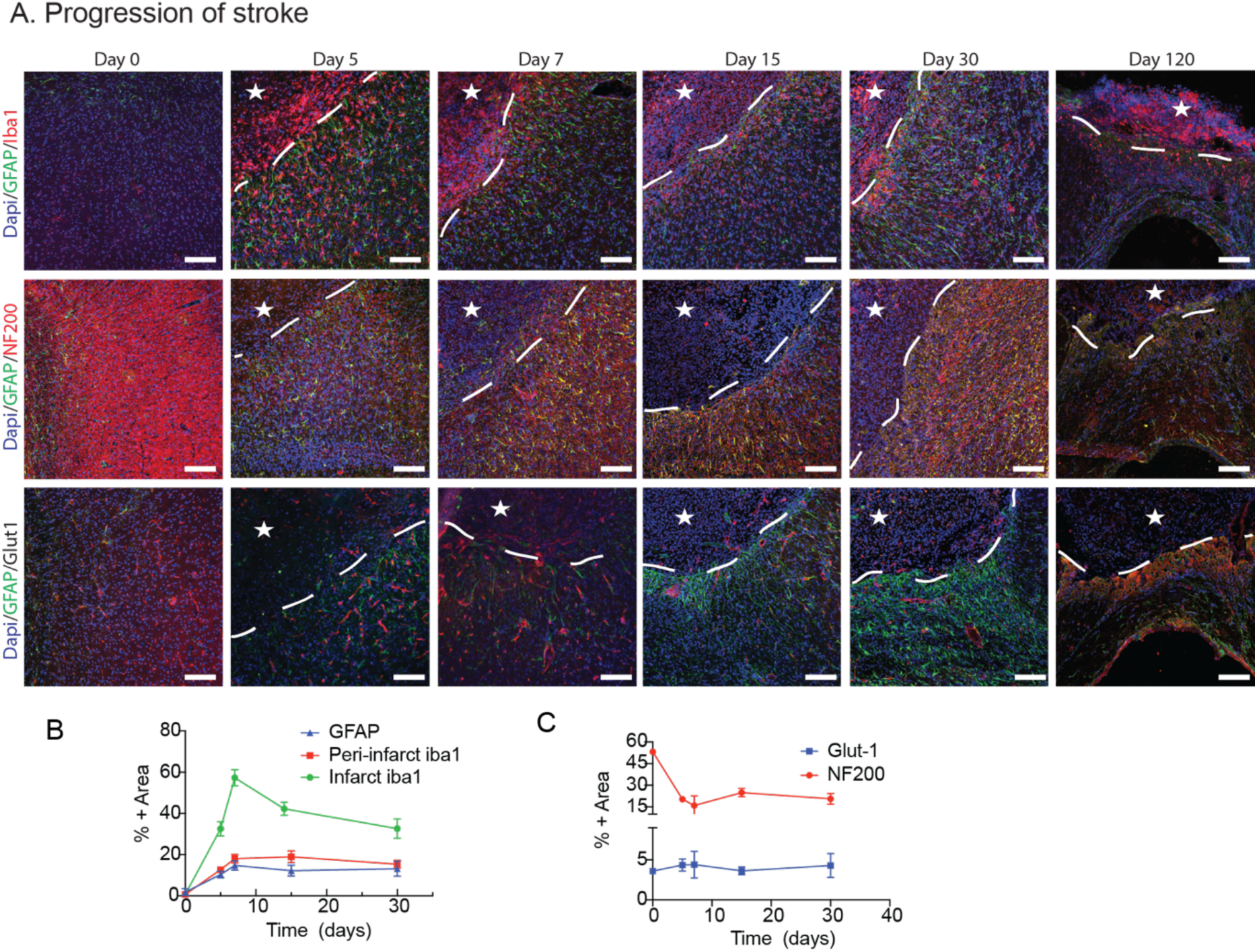
A. IHC images showing progression of inflammation (astrocytes and microglia/macrophages through GFAP and IBA-1 staining), axons (NF200), and vasculature (Glut1) beginning with healthy tissue and progressing to 5, 7, 15, 30, and 120 days after stroke. All scale bars = 100µm. B. Quantification of % area of inflammation over time through GFAP and IBA-1 staining. C. Quantification of % area over time of axons (NF200) and vessels (Glut1).

**Supplemental Figure 2:**
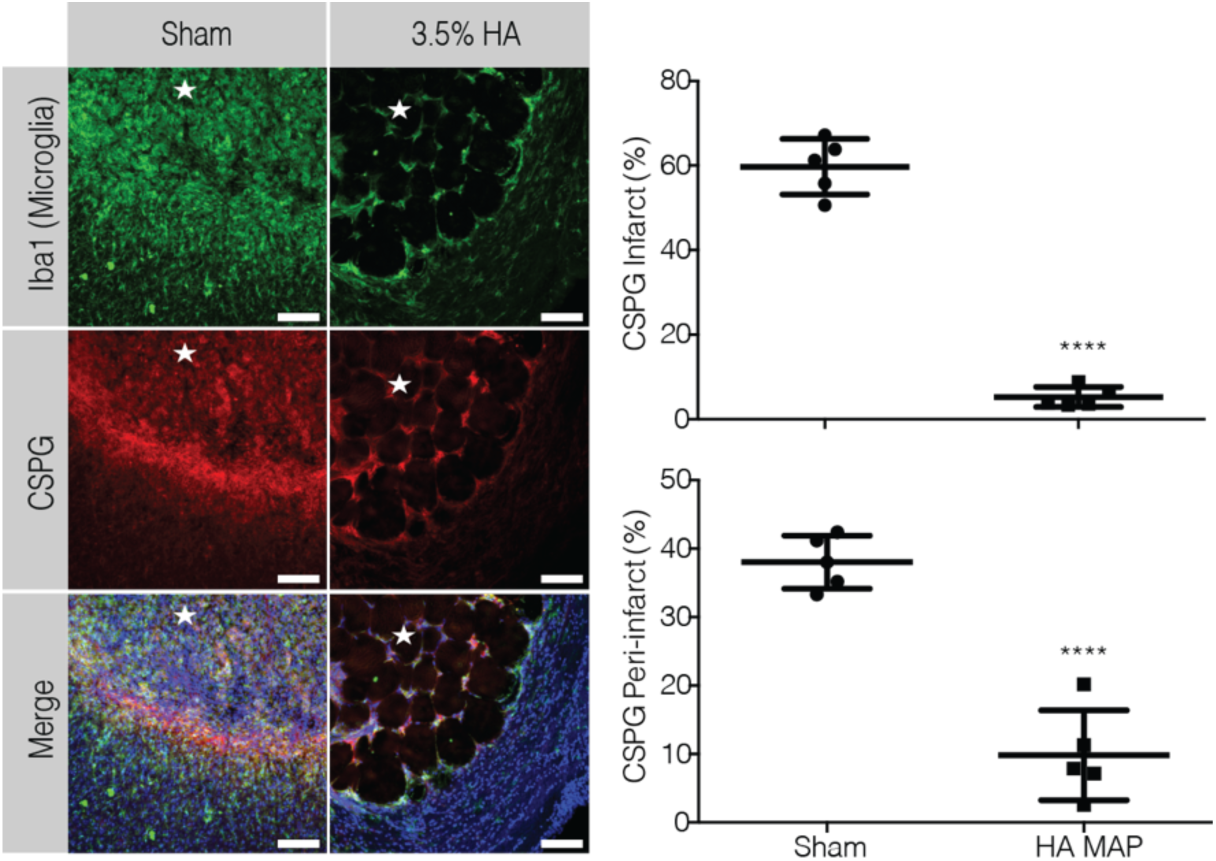
IHC images showing microglia and CSPG deposition at 7 day post stroke. Quantification of positive area of CSPG in peri-infarct and infarct comparing sham to MAP gel. All scale bars = 100µm. Statistical analysis was done in GraphPad Prism. Data were analyzed using a T-test and P<0.05.

**Supplemental Figure 3.**
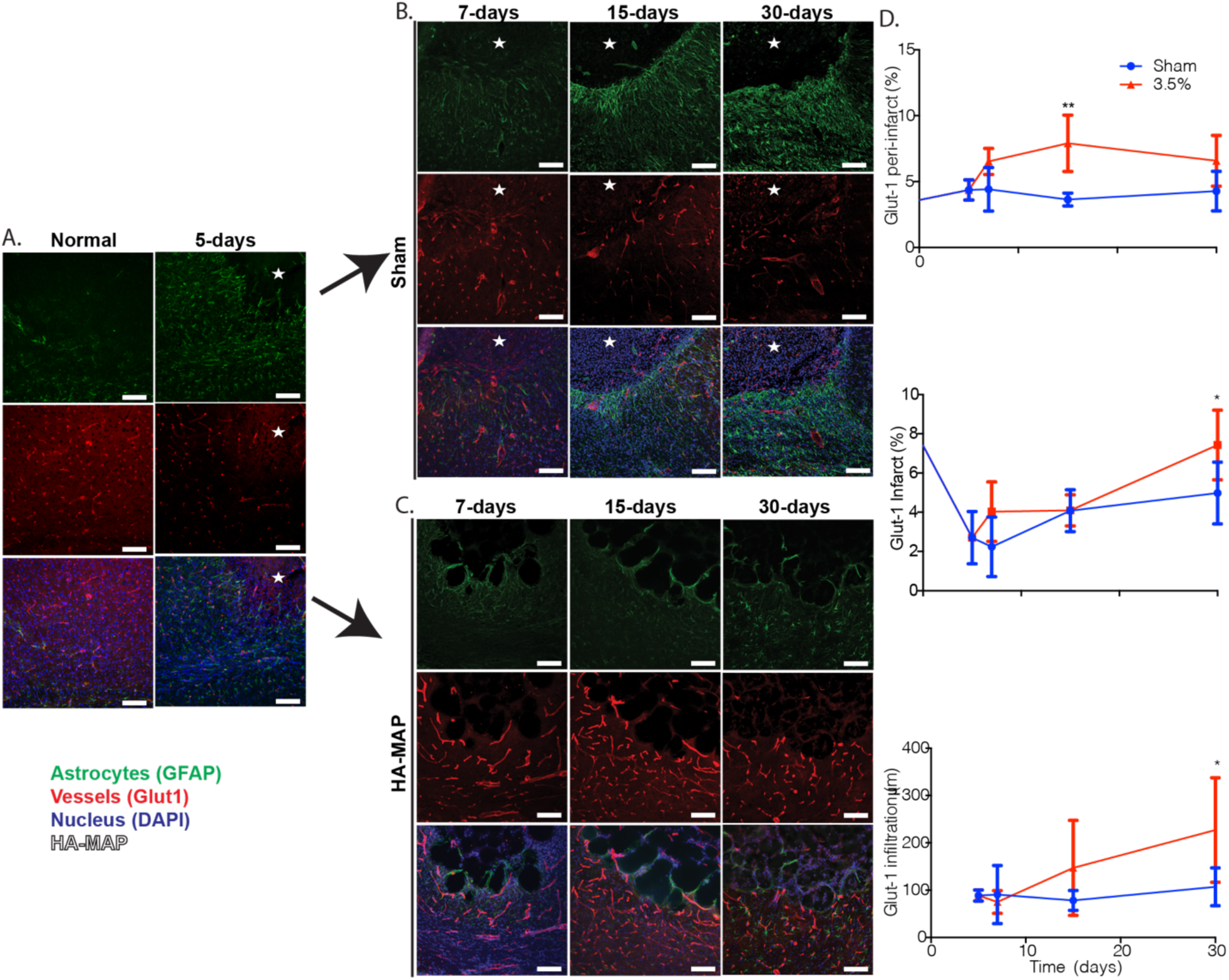
A. IHC images showing reactive astrocytes (GFAP) and vessels (Glut1) of healthy tissue and 5 days post stroke. B. IHC images of Sham stroke tissue showing reactive astrocytes (GFAP) and vessels (Glut1) at 7, 15, and 30 days post stroke. C. IHC images of MAP gel treated stroke tissue showing reactive astrocytes (GFAP) and vessels (Glut1) at 7,15, and 30 days post stroke, D. Quantification of percent area of vessels (Glut1) in the peri-infarct, infarct, and infiltration distance into infarct. All scale bars = 100µm. Statistical analysis was done in GraphPad Prism. Data were analyzed using a T-test and P<0.05.

**Supplemental Figure 4.**
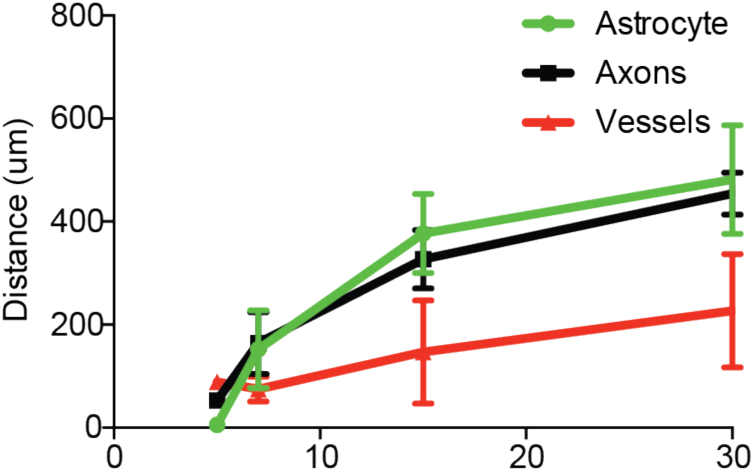
Overlay of infiltration of astrocytes (GFAP), axons (NF200), and vessels (Glut1) intro the infarct over time (7,15, and 30 days post stroke).

**Supplemental Figure 5.**
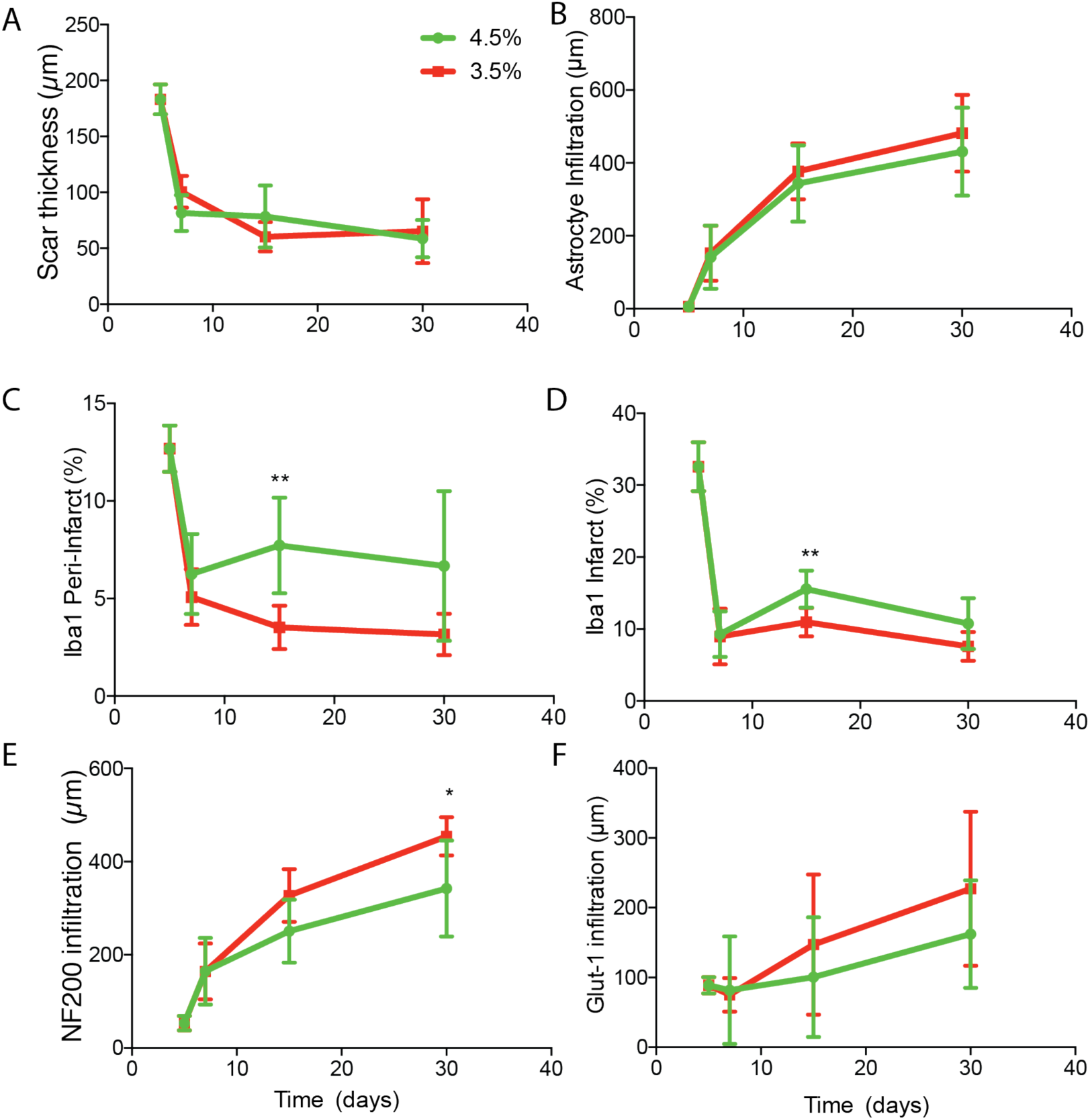
A. Quantification of scar thickness (GFAP) over time comparing 4.5% MAP and 3.5% MAP. B. Quantification of astrocyte infiltration (GFAP) into the infarct over time comparing 4.5% MAP and 3.5% MAP. C. Quantification of reactive microglia (IBA-1) percent area in the peri-infarct over time comparing 4.5% MAP and 3.5% MAP. D. Quantification of reactive microglia (IBA-1) percent area in the infarct over time comparing 4.5% MAP and 3.5% MAP. E. Quantification of axon (NF200) infiltration into the infarct over time comparing 4.5% MAP and 3.5% MAP. F. Quantification of vessel (Glut1) infiltration into the infarct over time comparing 4.5% MAP and 3.5% MAP. Statistical analysis was done in GraphPad Prism. Data were analyzed using a T-test and P<0.05.

**Supplemental Figure 6.**
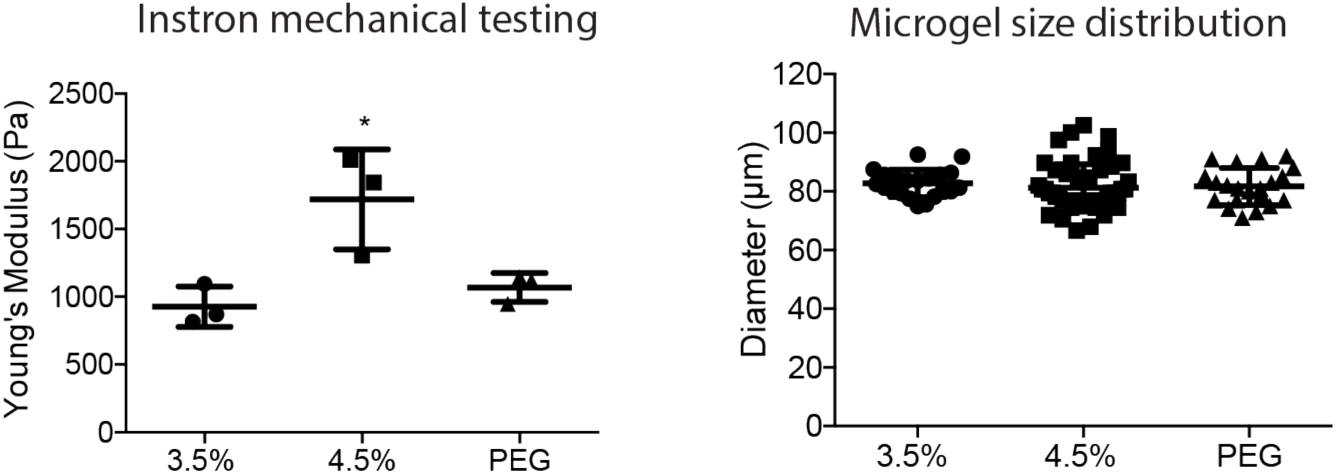
Quantification of Young’s modulus for 3.5% HA MAP, 4.5% HA MAP, and PEG MAP. Microgel size distribution for 3.5% HA MAP, 4.5% HA MAP, and PEG MAP. Statistical analysis was done in GraphPad Prism. Data were analyzed using a one way analysis of variance followed by a Tukey *post-hoc* test and a 95% confidence interval. * indicate P<0.05.

